# The phylogenetic Janzen-Connell effect can explain multiple macroecological and macroevolutionary patterns

**DOI:** 10.1101/2025.11.07.687215

**Authors:** Liang Xu, Hanno Hildenbrandt, Rampal S. Etienne

## Abstract

The classical Janzen-Connell (J-C) hypothesis states that species-specific natural enemies induce local-density dependence which explains high diversity observed in tropical tree communities. Recent studies have extended the hypothesis to include the fact that these natural enemies often attack phylogenetically related species as well, which reduces diversity and hence the explanatory power of the hypothesis. However, these studies consider ecological time scales at which the phylogenetic relatedness between species is constant, and thus ignore that at evolutionary time scales the population dynamics change the phylogeny. We use a spatially explicit eco-evolutionary model of the interplay between the dynamics of the phylogeny and the species abundances to study the predictions for common macroevolutionary and macroecological patterns. We confirm that hyperdiversity can be maintained by the J-C effect but the phylogenetic relatedness effect weakens the rarity advantage, reducing diversity. Incorporating protracted speciation further improves the result. Our model predicts a triphasic species-area relationship at much shorter time scales than in neutral scenarios. The species-abundance distribution can have one or two modes depending on the dispersal distance. Phylogenetic trees show diversification slowdowns and imbalance, consistent with empirical patterns. We study a new pattern of phylogenetic relatedness through space and find that small dispersal distance causes clusters of species with large phylogenetic distance to the community while large dispersal distance makes species distribute uniformly. As an illustration of how our model’s predictions can be compared to empirical data we study macroecological and phylogenetic patterns of the Barro Colorado Island (BCI) tree community. We conclude that the spatially explicit eco-evolutionary phylogenetic J-C effect can explain commonly observed macroevolutionary and macroecological patterns, providing an alternative way, while the traditional methods are prone, to demonstrate the role of J-C effect on species community assembly.

## INTRODUCTION

The high diversity in many ecological communities, such as the Amazon rain forest hosting approximately 16,000 tree species (Ter Steege et al. 2013), never ceases to amaze. How such diversity is maintained has puzzled biologists for decades. One of the leading explanations of high biodiversity is the Janzen-Connell (J-C) effect (Janzen 1970; Connell 1971), which posits a reduction of species recruitment in areas of high conspecific density, because natural enemies such as seed predators, parasites, herbivores and pathogens are attracted by locally abundant species. As a result, rare species are at a recruitment advantage, leading to high diversity. The J-C effect thus involves density-dependence in a spatial context where the attenuation of the effect with distance has been referred to as distance-dependence.

Since the J-C hypothesis was first proposed, it has received much attention. The number of articles on the J-C hypothesis is substantial (Comita et al. 2014) and review papers that summarize studies testing the J-C hypothesis have been periodically published (Clark and Clark 1984; Hammond and Brown 1998; Takeuchi and Nakashizuka 2007). In a meta-analysis of empirical studies, Comita *et al*. (Comita et al. 2014) reported strong support for both distance- and density-dependent predictions. Evidence has also accumulated that natural enemies are likely to attack multiple, closely related, species (Novotny et al. 2002; Gilbert and Webb 2007; Liu et al. 2012). If this phylogenetic J-C effect attenuates with phylogenetic distance, its ability to explain high diversity may be compromised, because rare species that have abundant phylogenetic relatives have a reduced advantage if their relatives are more common (Gilbert and Webb 2007; Ness et al. 2011). However, the theory of phylogenetic J-C effect has not been studied yet.

It has been recently argued that the strength of the Janzen-Connell effect is likely to be overestimated using direct field measurements (Detto et al. 2019), and may therefore be falsely detected when actually absent. An alternative and complementary approach to find support for the J-C effect is to study its effect on macroecological and macroevolutionary patterns. Various studies have therefore modelled the J-C effect and explored the role of phylogenetic relatedness (Muller-Landau and Adler 2007; Chisholm and Muller-Landau 2011; Mack and Bever 2014; Miranda et al. 2015; Stump and Chesson 2015; Stump and Comita 2018) on these patterns. For example, Muller-Landau & Adler (Muller-Landau and Adler 2007) demonstrated that large dispersal distance of natural enemies increases species richness while plant dispersal has a complex effect on diversity. Miranda *et al*. (Miranda et al. 2015) showed that strong J-C effects lead to high diversity and that a small variance in the negative density dependence among species can explain high diversity. Levi *et al*. (Levi et al. 2018) incorporated distance-responsive natural enemies into the classic neutral theory (Hubbell 2001) to account for the J-C effect and confirmed that the J-C mechanism can lead to hyperdiversity. However, they did not consider the phylogenetic relatedness effect and may thus have overestimated species richness. Sedio & Ostling (Sedio and Ostling 2013) did model the phylogenetic J-C effect in a spatially explicit model to study how specialized natural enemies need to be to promote coexistence. They only considered ecological time scales and hence set a fixed phylogenetic distance among species. On (macro)evolutionary time scales the phylogenetic relationships are continuously changing, as they are affected by the J-C dynamics themselves. The consequences of this feedback for macroeological and macroevolutionary patterns have not been studied.

Here, we explicitly model this feedback and integrate it with all other previously studied factors by developing an individual-based dynamic eco-evolutionary model with a spatially explicit phylogenetic J-C effect. Our model is a non-neutral extension of the standard neutral model of biodiversity in which the continuously changing phylogeny and abundance affect recruitment in a spatial grid. The model has four key parameters, i.e. the strength of the Janzen-Connell effect (i.e. how disproportionately natural enemies are attracted by abundant species), the strength of phylogenetic relatedness (i.e. how quickly the J-C affect attenuates with phylogenetic distance), the interaction distance of the J-C effect (i.e. how quickly the J-C effect attenuates with spatial distance) and the dispersal distance of individual plants. We explore the predictions of our spatial phylogenetic Janzen-Connell (SPJC) model for species richness, the species abundance distribution (SAD), the species-area relationship (SAR), phylogenetic properties such as tree balance and lineages-through-time (LTT) plots, and a new metric for the phylogenetic distribution in space. We confirm that hyperdiversity only results when pathogens or predators are very species-specific. Furthermore we find that the spatially explicit phylogenetic J-C effect can generate a variety of ecological and phylogenetic community patterns. We provide a new spatio-phylogenetic prediction that small dispersal causes the formation of spatial clusters of species with large phylogenetic distance to the community whereas for large dispersal distance of individual plants phylogenetic relatedness is evenly distributed across space. We compare our model’s predictions qualitatively with tree community data from Barro Colorado Island, and find support for a strong phylogenetic Janzen-Connell effect and intermediate dispersal limitation. However, we do not want to draw firm conclusions because the spatial scale of the BCI community (local) does not match the scale of our simulations (regional); the empirical example mostly serves as an illustration. We argue that our model provides a mechanistic explanation for diversity-dependent diversification, and that adding further model details such as time-dependence and age-dependence of the phylogenetic Janzen-Connell and increasing grid size can improve our match with empirical data.

## MODEL DESCRIPTION AND ANALYSIS

### The spatially explicit phylogenetic Janzen-Connell model and simulation process

We consider species dynamics in a spatial grid of *N* cells, each cell hosts one individual. We assume that at each time step one individual dies, chosen at random, as in the standard, non-spatial, neutral model (Hubbell 2001; Etienne 2005) as well as spatially explicit versions (Rosindell and Cornell 2007; Rosindell et al. 2008). The empty cell is colonized by offspring of an individual anywhere in the grid, depending on its proximity. However, in contrast to the neutral model, the probabilities of colonizing the empty cell are not equal for all individuals at the same spatial distance, but depend on the phylogenetic distance to the individuals surrounding the empty cell (and hence this also accounts for their spatial distance and - at the species level - the abundance). We assume that the colonization probability of (the offspring of the) individual in cell *k* belonging to species *i* to the empty site *o* is given by

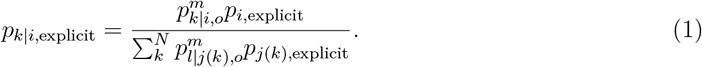

The formula (Eq. 1) contains two components, i.e. the dispersal probability 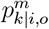 of individuals from cell *k* to the empty cell *o* and the relative probability of species *i* colonizing the cell that accounts for the phylogenetic J-C effect. The relative probability of species *i* colonizing the cell *o* is assumed as

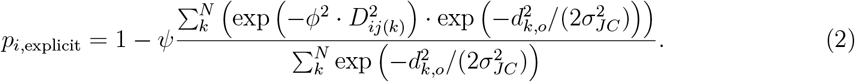

where *D*_*ij*(*k*)_ is the phylogenetic distance between species *i* and the species identity (for example, species *j*) of the individual in cell *k, d*_*k,o*_ is the distance between cell *k* and the empty cell *o, ψ* determines the strength of the J-C effect, *σ*_*JC*_ determines the spatial extent of the J-C effect, and *ϕ* measures the size of the phylogenetic effect. The Gaussian kernel exp 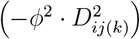 describes the differential contribution to the colonization probability from very small for a phylogenetically remotely related species (large *D*_*ij*(*k*)_) to very large for a conspecific (*D*_*ij*(*k*)_= 0) . When the J-C effect is absent, i.e. *ψ* = 0, the model reduces to a spatially explicit neutral model. When *ϕ* approaches infinity, the J-C effect is restricted to closely related species as in the classic J-C hypothesis (only the term *D*_*jj*_ contributes to the sum), but when *ϕ* is small, even a species *i* that is phylogenetically distant from the individuals surrounding the empty cell experiences a reduced colonization probability; when *ϕ* = 0, we arrive again at the neutral model, because all species are affected equally (they all have the same relative probability). When *σ*_*JC*_ is small, only individuals immediately surrounding the empty cell determine the colonization probability (i.e. those individuals attract enemies that attack their conspecifics and phylognetically related species and thus reduce these species’ colonization probability).

The phylogenetic distance between all species pairs can be summarized in a matrix **D**, defined by

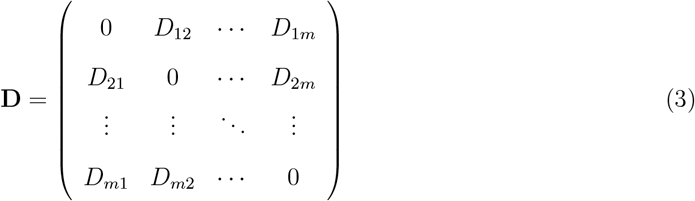

where the entry *D*_*ij*_ denotes the interspecific phylogenetic distance between species *i* and *j*. We thus do not model the dynamics of the host-pathogen interaction explicitly, but use the phylogenetic distance as a proxy for the likely host range of natural enemies.

The probability of dispersal 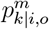 of the individual in cell *k* belonging to species *i* to the vacant site *o* is weighing the relative specie colonization probability by considering the geographic distance. We assume that this probability follows a normal distribution determined by the geographic distance from the individual’s site to the vacant site, *d*_*k,o*_, and the dispersal distance of individuals *σ*_*disp*_ (Rosindell and Cornell 2012), which is assumed to be the same across all species:

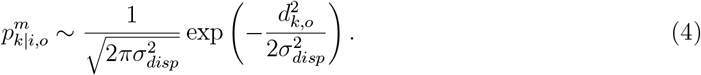

When *σ*_*disp*_ is infinite, all individuals can move anywhere in the community with equal probability. Other dispersal kernels such as the fat tailed kernel (Rosindell and Cornell 2007, 2012) can also be implemented, but we note that the dispersal kernel does not affect the pattern (Rosindell and Cornell 2007) much. For this reason, and because the analysis of our simulation scenarios are very computationally expensive we have not explored predictions for alternative dispersal kernels.

The model reduces to a non-spatial model when setting the dispersal ability of individuals *σ*_*disp*_ and the spatial interaction distance of the phylogenetic J-C effect *σ*_*JC*_ to infinity. Then we obtain the non-spatial colonization probability of an individual of species *i*

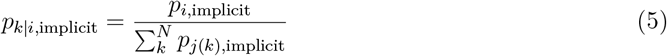

where the non-spatial relative colonization probability of species *i* is given by

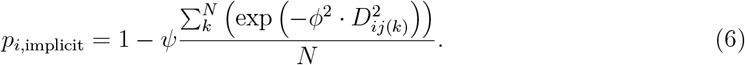

We illustrate how the phylogenetic J-C effect affect the colonization probability by comparing the relative colonization probabilities (Eq. 6) for different *ϕ* (Fig.1) with a simple tree of 4 species and their abundances in a non-spatial context .

**Figure 1.**
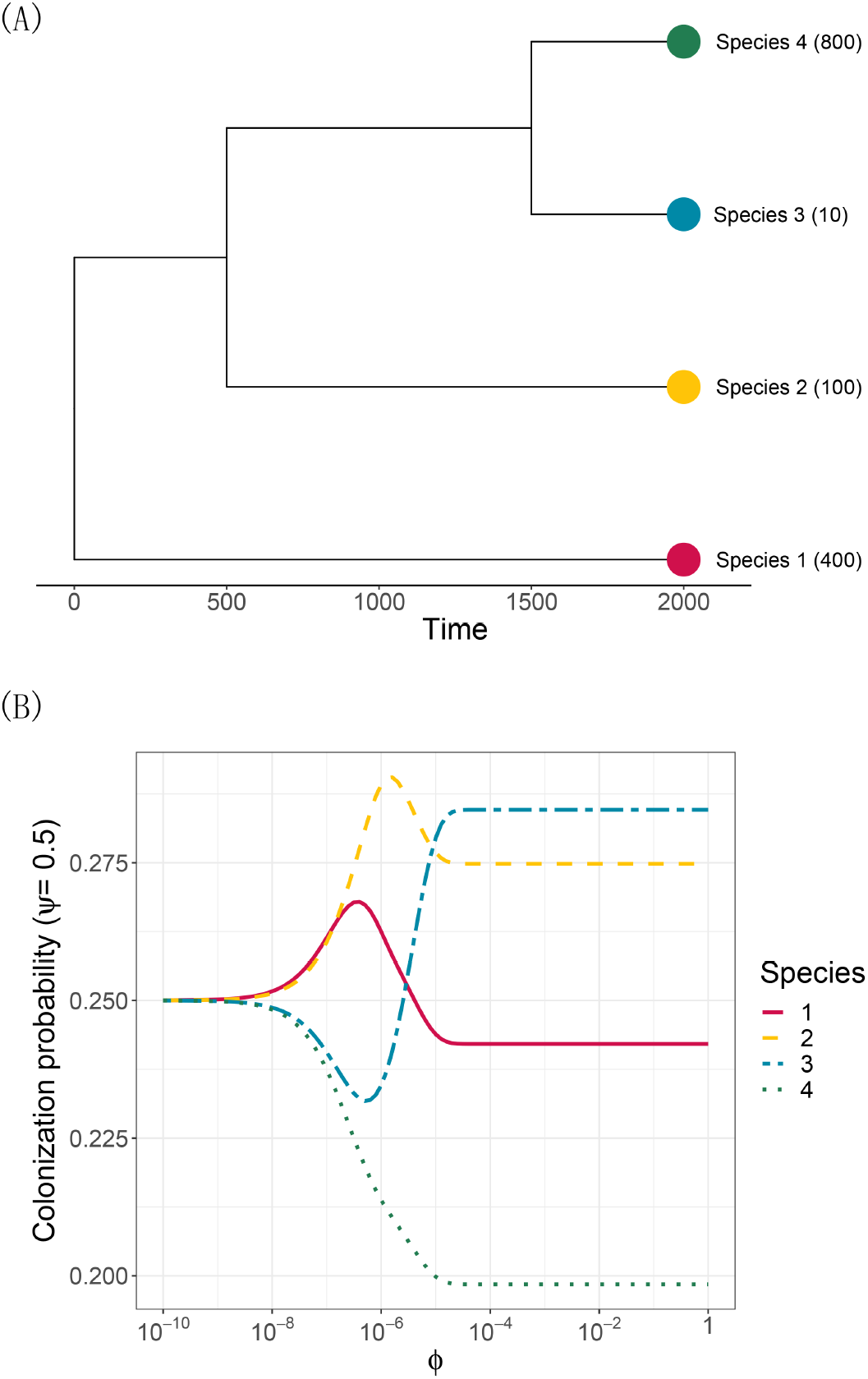
The relative colonization probabilities for 4 species of a simple tree when *ψ* = 0.5. Panel A shows the tree with tips labeled by their abundances. The phylogenetic relatedness matrix is therefore given by 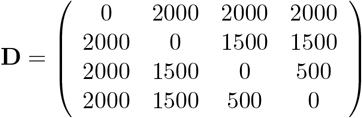. Panel B shows the colonization probabilities against the variable *ϕ*. When *ϕ* is small, the rare species (Species 3) suffers the negative effect from phylogenetically related species, thus, possesses a low colonization probability as Species 4. With increasing *ϕ*, natural enemies become more host-specific and the rarity advantage increases due to the J-C effect. Thus, Species 3 is less affected by its conspecific density and finally reaches a highest colonization probability.

We assume that the community has a fixed size *N* and we start with two species, one common species with abundance *n*_1_ = *N* − 1 and one rare species with abundance *n*_2_ = 1. The initial phylogeny tree has two equivalent branches. The initial phylogenetic distance matrix **D** can be written as 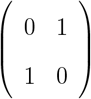. At each time step (Fig. 2), one individual in the community is randomly chosen to be removed (Levi et al. 2018). The abundance of the corresponding species decreases by one. If the species is a singleton, the death event results in extinction. We then remove that species from the community and the phylogeny. The corresponding column and row in the phylogenetic distance matrix **D** are removed as well. After random death the cell is colonized immediately. The colonizer is a new species if speciation occurs with probability *v* or offspring of an existing species with probability 1 − *v*. If birth occurs, one individual is sampled according to the colonization probability (Eq. 1). After the birth event, the abundance of the corresponding species increases by 1. If speciation occurs, we sample one individual again using the colonization probability (Eq. 1) and let the corresponding species speciate. To update the abundance and phylogenetic relatedness, we extend the abundance vector by adding the new species with 1 individual (the point mutation assumption Hubbell 2001) and we extend the phylogenetic relatedness matrix by adding one column and one row. Because the point mutation assumption often generates an unrealistically large number of very rare species (Rosindell et al. 2010), we exploited the idea of the protracted speciation model that considers that speciation takes time (Rosindell et al. 2010). We introduced the completion time *τ* (we studied *τ* = 0, 10^4^, 10^5^, 5 *×* 10^5^, 10^6^ where *τ* = 0 represents the point mutation speciation model) for the newly born species such that these species are identified as species only when they are still present after *τ* generations. Therefore, only the extant species that were produced before *τ* generations are considered. The phylogenetic relatedness of the new species to the others are the same as that to the sampled parent species (plus 1) while to the parent species this distance equals 1. Finally, at every time step we update the phylogenetic relatedness matrix (Eq. 3) by adding 1 to every off-diagonal entry. These steps are repeated until a prescribed time is reached. We call this sequence of events the tree replacement. The model is coded in C++ and the analysis is in R. All the code can be found on …

**Figure 2.**
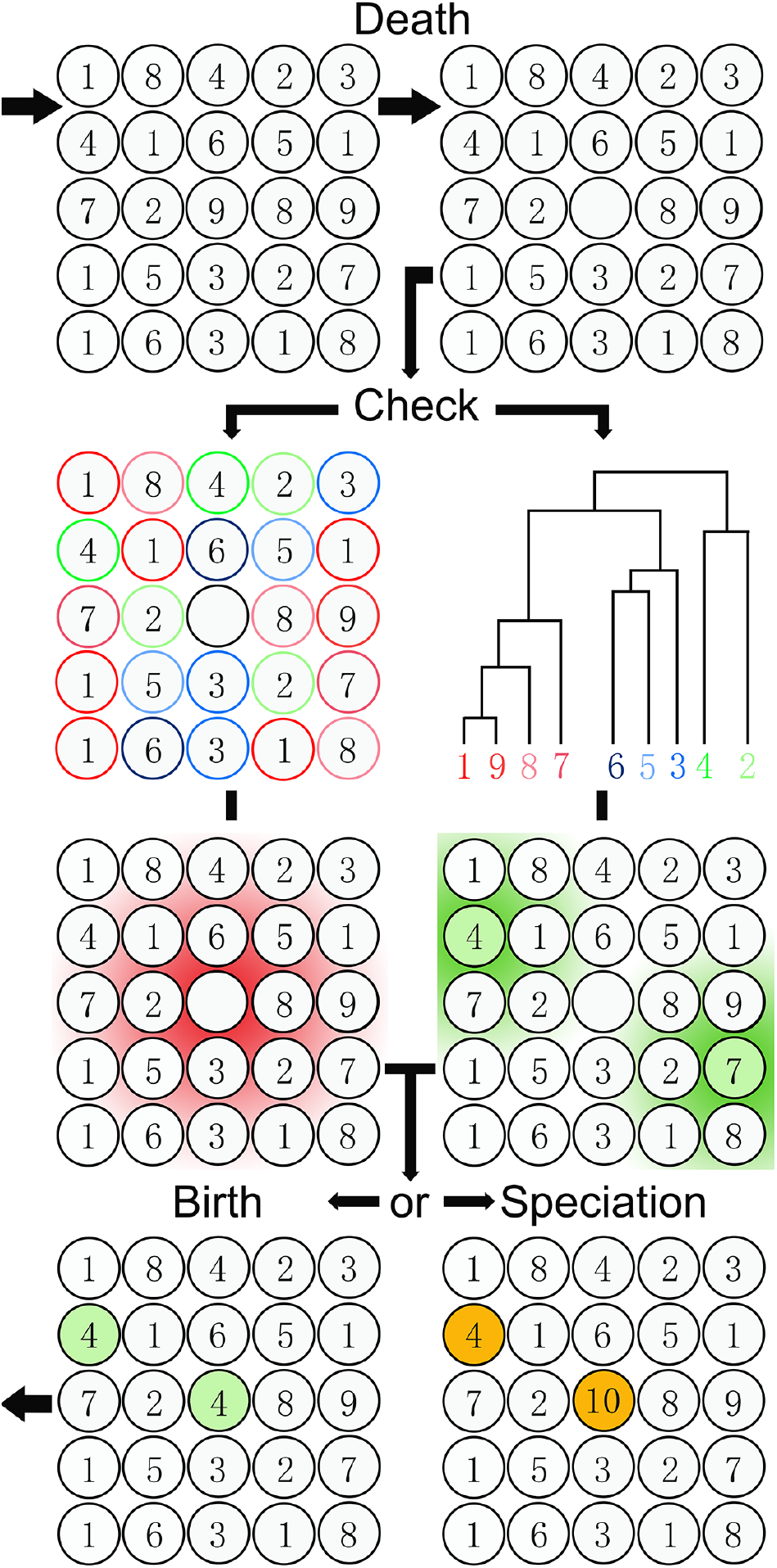
Schematic representation of one tree replacement of the model for a grid size of *N* = 5^2^. Each cell can host one tree which is labeled with a number. After one individual’s death, the individual colonization probability is computed for all individuals based on their phylogenetic relatedness (given by the phylogenetic tree) and density (denoted by the number of circles with the same number and color) and further weighted by the interaction distance of the phylogenetic J-C effect (denoted by the red zone around the vacant site in the center) and the dispersal distance of individuals (the green zone around individuals; dispersal zones of only two individuals are shown). The chosen individual gives birth to a new individual, but this individual may undergo speciation with probability *ν* = 0.0001 (species 10 arises from speciation in species 4) and inherits the phylogenetic distance to other species from its parent.

### Parameter settings

Our aim is to examine to what extent the phylogenetic J-C effect results in different patterns compared to neutral theory and how different J-C interaction distances and dispersal distances affect these patterns, so we can compare these predictions to empirical patterns (see below). To assess the effect of different spatial scales on diversity and better compare with the empirical data that are on local community scales, we set the regional grid size to *N* = 333^2^ and studied the diversity of two additional local scales within this grid (*N* = 55^2^, 111^2^) by sampling all possible non-overlapping areas of that scale in the grid and measure the species richness. Additionally we subsampled communities of size 50 *×* 50, 100 *×* 100, 150 *×* 150, 200 *×* 200, 250 *×* 250 in the middle of the grid of each simulation and took the average across 100 replicates for each parameter setting to study various macroecological and macroevolutionary patterns (see below) at these sampling scales. We varied the strength of the phylogenetic J-C effect *ψ* in the range (0, 0.25, 0.5, 0.75, 1), which describes scenarios ranging from neutrality to a very strong J-C effect. The parameter *ϕ* quantifies the width of the phylogenetic relatedness effect which we vary in the range of (1, 10^−2^, 10^−4^, 10^−6^, 10^−8^, 0). Note that *ϕ* = 1 approaches the scenario of the classic J-C effect and *ϕ* = 0 refers to the neutral scenario. The speciation rate is set at *ν* = 0.0001,which means that there are 10^4^ tree replacements on average between speciation events. The total number of tree replacements in the simulation (which determines the maximum phylogenetic distance) was set to 10^7^ for computational tractability. We chose three values each for *σ*_*disp*_ and *σ*_*JC*_: (0.1, 1, 10) referring to low distance, intermediate distance and high distance, which results in 9 distance scenarios. We ran 100 replicate simulations for each of the 270 parameter combinations leading to a total of 27000 simulations, each of which took up to 5 days.

### Model analysis

We studied the predictions of our model for the species abundance distribution (SAD), the species-area relationship (SAR), the lineages-through-time (LTT) plot, the phylogenetic balance and a new metric that shows how the phylogenetic J-C effect affects the spatial distribution of the species. This new metric is based on the mean of inverse pairwise phylogenetic distances (MIPD, Ness et al. 2011). In contrast to other widely used metrics such as the phylogenetic distance to the phylogenetically nearest neighbor (NND, Webb et al. 2006) and mean phylogenetic distance to all other species (MPD, Vamosi et al. 2009; Hill and Kotanen 2009), MIPD increases with increasing relatedness of the focal species to others in the community nonlinearly, thus measuring species-level phylogenetic similarity in a community. However, the mean of inverse distances overweights the influence of phylogenetically closely related species and does not take into account abundance. Therefore, for our metric of phylogenetic similarity we take the exponential rather than the inverse and weigh the result with abundances: the abundance-weighted MIPD, or AWMIPD. For species *i*, AWMIPD_*i*_ is defined as:

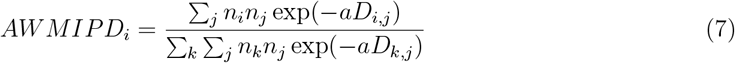

where *n*_*i*_ is the abundance of species *i, a* is a scalar that sets the scale of the effect of phylogenetic distance. A species can attain a high AWMIPD if it is closely related to the other species (small *D*_*i,j*_), or if either species has a high abundance (i.e. large *n*_*i*_ or *n*_*j*_). The scalar *a* can be chosen by the user; it depends on the range and units of the phylogenetic distances. We tested *a* = 0.01, 0.1, 1, 10, 100, 1000 for phylogenies with normalized crown age (i.e. crown age set to 1). There are no large differences among them (Fig. S1 - S6), so we chose *a* = 1 for all the AWMIPD calculations. We plotted the AWMIPD values for all individuals across space.

To analyze the phylogenetic tree balance, we computed the Colless index *I*_*m*_ (Colless 1982; Mooers 1995; Bortolussi et al. 2006) and the *β*-statistic (Aldous 1996, 2001; Blum and François 2006; Bortolussi et al. 2006). The Colless index *I*_*m*_ ranges from 0 (balanced trees) to 1 (imbalanced trees). For the *β*-statistic, *β* = 0, corresponds to the balance of a pure-birth process (the Yule model Yule G. 1925). A positive *β* value indicates a more balanced tree than the Yule process while a negative value implies a more unbalanced tree.

We measured the rate of lineage accumulation with the *γ*-statistic (Pybus and Harvey 2000) and the Δ*r*-statistic (Pigot et al. 2010; Etienne and Rosindell 2012). The *γ*-statistic is a commonly used index of species accumulation but it depends on the size of the tree (McPeek 2008; Etienne and Rosindell 2012). When *γ >* 0, the phylogeny’s internal nodes are closer to the tips than to the root, which indicates an accelerating pattern in speciation. Otherwise when *γ <* 0, the phylogeny shows a slowdown in lineage accumulation. The Δ*r*-statistic is an alternative statistic which is computed as (Magallón and Sanderson 2001; Pigot et al. 2010)

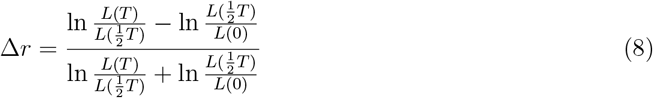

where *L*(*t*) denotes the number of lineages at time *t* that survive up to the present and *T* is the crown age, i.e. the time since the most recent common ancestor of all species in the community at the present. The parameter Δ*r* varies from −1 to 1, with Δ*r* = 0 indicating constant lineage accumulation, Δ*r >* 0 indicating acceleration in, and Δ*r <* 0 indicating a slowdown in lineage accumulation.

We compared our model’s predictions on different subsampling scales to empirical data of the Barro Colorado Island (BCI) tree community to explore if our model could generate realistic eco-evolutionary patterns. The BCI data set consists of the locations of all woody trees and shrubs with stems at least 1 cm in stem diameter in a 50 ha plot (500 m *×* 1000 m) (Condit 1998; Hubbell et al. 1999, 2005; Condit et al. 2012; Pearse et al. 2013). Because the phylogenetic relatedness effect and the Janzen-Connell effect may have distinct effects on different life stages (Zhu et al. 2015) we filtered plants with three diameters at breast height (dbh > 10 mm, 30 mm, 100 mm) and analyzed the patterns for each. Although the community size of this data set may be not sufficiently large to show substantial macroevolutionary patterns (see Discussion), we used it anyway as an example to illustrate how to compare and analyze the simulated patterns with the observations. The BCI data are available as “Barro Colorado Forest Census Plot Data (Version 2012)” at https://doi.org/10.5479/data.bci.20130603, Condit et al. 2012.

Furthermore, to compare our generated phylogenetic metrics to observations more generally, we used two data sets, each with many phylogenetic trees (from hereon referred to as ‘phylogenetic data sets’): one from TreeBASE (www.treebase.org/) and one used by McPeek (McPeek 2008). After filtering out the trees that lacked information on branch lengths, or had less than 3 extant species (Davies et al. 2011), we finally arrived at 573 and 145 phylogenetic trees, respectively, for comparing the metrics to our simulated data. We note that these empirical phylogenies cover a wide range of taxa, including taxa that may not be associated with a J-C effect. We used them anyway to get an idea of the range of empirical phylogenetic patterns.

## Results

### Species richness

The J-C effect promotes biodiversity but less so when the J-C effect extends to phylogenetically more remote species (small *ϕ*). When *ψ* = 0 or *ϕ* = 0, the model reduces to the (spatially explicit) neutral model. We will therefore call the combinations with *ϕ* = 0 or *ψ* = 0 the neutral parameter combinations, and refer to other parameter sets as the non-neutral parameter combinations. For nonzero *ψ* species richness increases with increasing *ϕ* (see Fig. 3). At the largest values of *ϕ*, the system resembles the classic J-C scenario which has highest diversity. Small dispersal distance produces more species than large dispersal distance for all parameter combinations at the regional scale (*N* = 333^2^). In contrast, larger interaction distance of the phylogenetic J-C effect results in more species than a smaller interaction distance, but the increase is not as substantial as that produced by changing the dispersal distance. Thus, the highest species richness is observed when the dispersal distance is smallest (*σ*_*disp*_ = 0.1) and the interaction distance of the phylogenetic J-C effect is largest (*σ*_*JC*_ = 10) together with the largest values of *ϕ* and *ψ* (*ϕ* = 1, *ψ* = 1). This species richness is almost 10 times larger than the richness that is generated by the standard neutral theory in the same grid. For small spatial scales (*N* = 55^2^, 111^2^, Fig. 3), we found that increasing dispersal distance increases species richness, implying that changing the scale from large to small shows a change in the dispersal-diversity relationship from negative to positive. Incorporating the protracted speciation model does not change the patterns across different dispersal distances. However, under the protracted speciation model the species richness decreases more for the neutral scenarios than the scenarios with the phylogenetic J-C effect (Fig. S52) under large *τ* values (*>* 10^5^).

**Figure 3.**
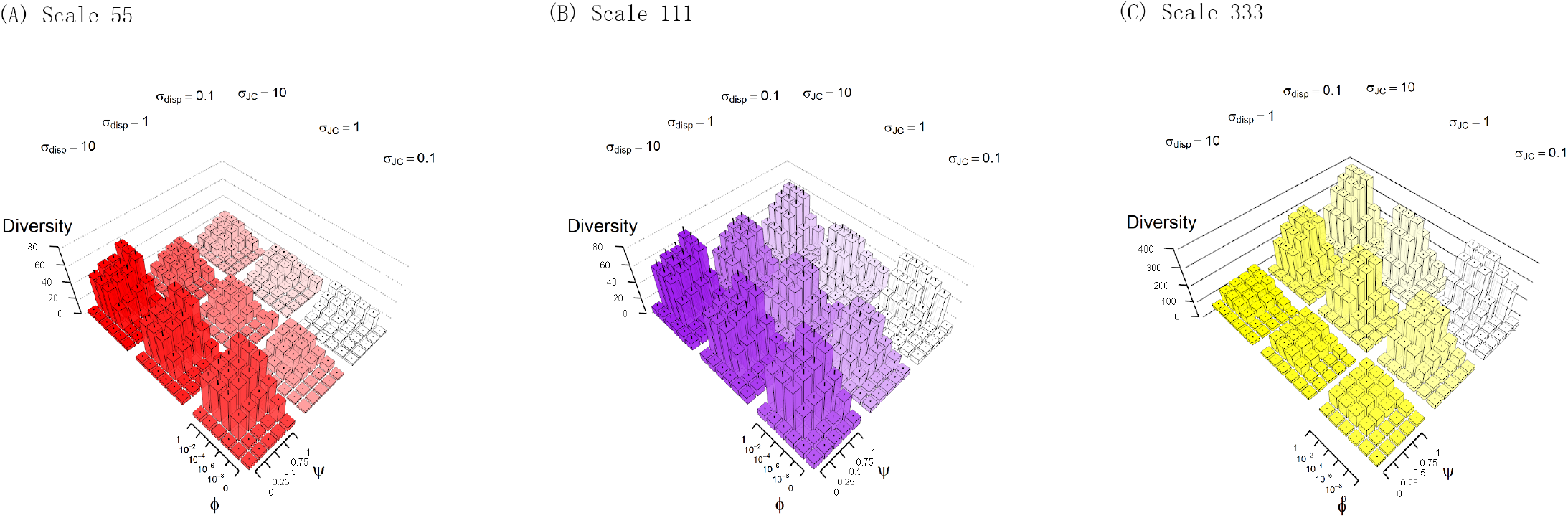
Species richness for increasing spatial scale (A-C). In each panel, the 9 blocks of 30 bars represent the combinations of three different interaction distances of the phylogenetic J-C effect and three different dispersal distances. Each block of 30 bars represents 30 combinations of *ϕ* and *ψ*. The top surface of each bar is the mean diversity across 100 replicates, and the error line in each bar represents the first and third quantiles. To study the species richness pattern on one specific subsampling scale, one can focus on the corresponding subplot. The 9 blocks in each subplot indicate the pattern for different interaction and dispersal distances. Further zooming in on each block, one can study the diversity pattern for different strengths of the phylogenetic J-C effects.

### Species-area relationships

The phylogenetic J-C effect generates a clear triphasic species-area relationship on the whole grid scale, which is in line with data and neutral model predictions (Hubbell 2001; Rosindell and Cornell 2007), for all parameter settings, except, remarkably, the neutral scenario (Fig. 5b and Figs. S7-S15). However, this is due to the fact that the neutral simulations have not reached equilibrium. Indeed, running the simulations longer results in a triphasic SAR for the neutral scenario as well (Fig. S16). Hence, the phylogenetic J-C effect creates the triphasic shape much faster than the neutral model can. On smaller sampling scales, the SAR tends to show the first two phases. There is little difference in the shape of the SAR between the scenarios with and without protracted speciation.

**Figure 4.**
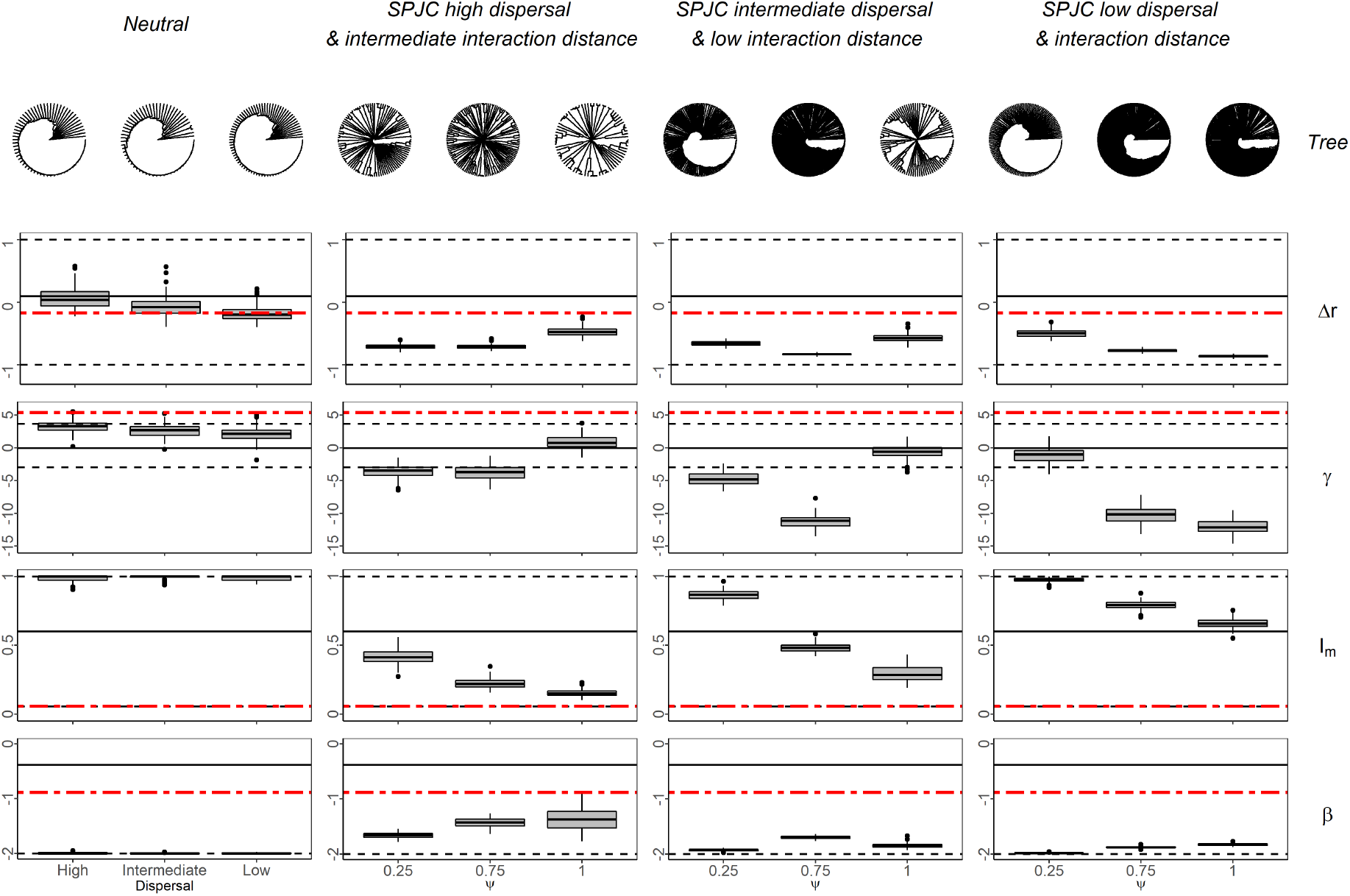
Distinct distributions of species accumulation and imbalance statistics produced by different parameter combinations. The generating parameters from left to right are (1) for the neutral scenarios (with *ψ* = 0, *σ*_*JC*_ = 10): *σ*_*disp*_ = 10; *σ*_*disp*_ = 1; *σ*_*disp*_ = 0.1, (2) for the large dispersal distance and intermediate interaction distance scenarios (*σ*_*disp*_ = 10; *σ*_*JC*_ = 1): *ψ* = 0.25, *ϕ* = 10^−4^; *ψ* = 0.75, *ϕ* = 10^−4^; *ψ* = 1, *ϕ* = 10^−6^, (3) for the intermediate dispersal distance and low interaction distance scenarios (*σ*_*disp*_ = 1; *σ*_*JC*_ = 0.1): *ψ* = 0.25, *ϕ* = 10^−4^; *ψ* = 0.75, *ϕ* = 10^−4^; *ψ* = 1, *ϕ* = 10^−6^, (4) for the small dispersal distance scenarios (*σ*_*disp*_ = *σ*_*JC*_ = 0.1): *ψ* = 0.25, *ϕ* = 10^−4^; *ψ* = 0.75, *ϕ* = 10^−4^; *ψ* = 1, *ϕ* = 10^−6^ respectively. The phylogenetic trees each represent one sample out of 100 replicates generated by the aforementioned parameters. In the box plots, solid lines, boxes and whiskers denote the 50th, 25th/75th and 5th/95th percentiles, respectively. The solid horizontal line and dashed horizontal lines represent the median and interquartile range (5%-95%) of empirical estimates of the corresponding statistic for 573 phylogenetic trees in TreeBASE, respectively. The red horizontal line represents the estimates of the corresponding statistic for the BCI data.

**Figure 5.**
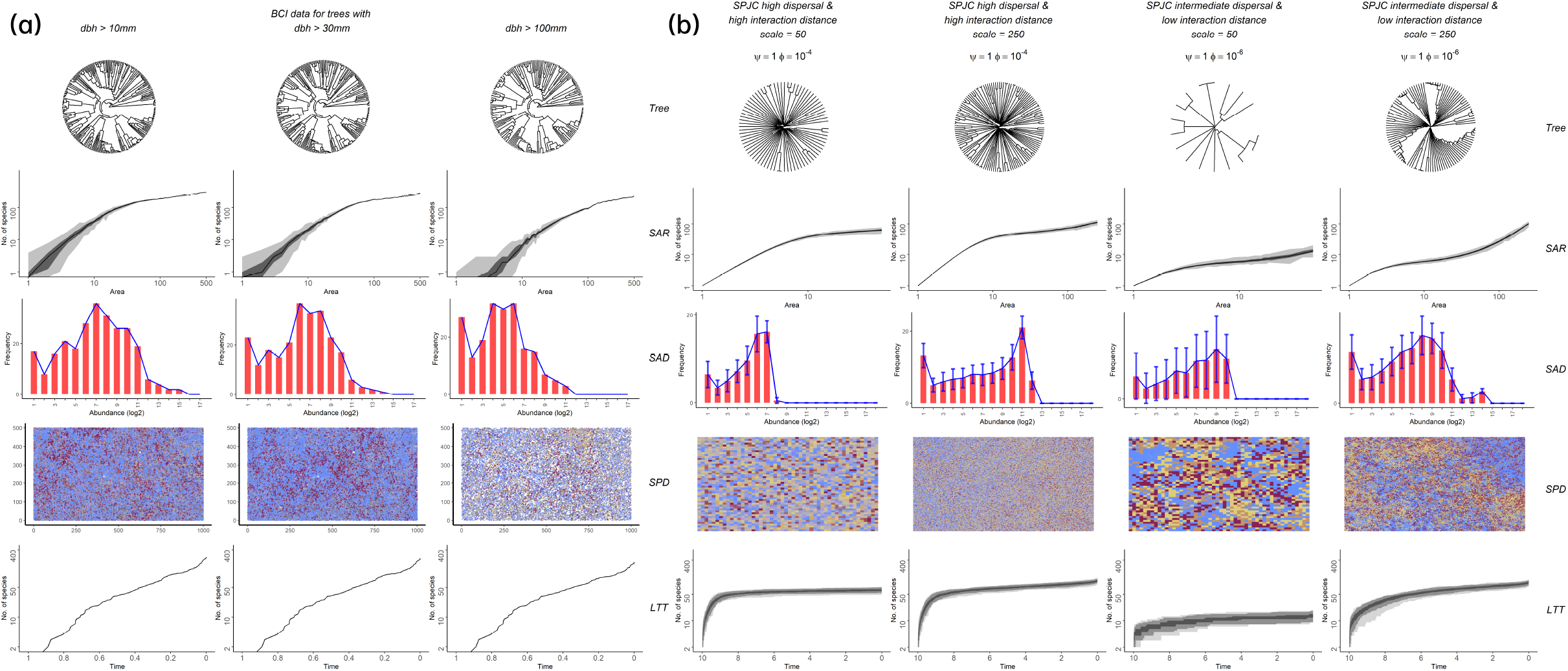
Distinct ecological and phylogenetic patterns of the BCI data on three dbh values, i.e. dbh > 10 mm, 30 mm, 100 mm, respectively (a) and of the different parameter combinations and on different spatial scales (b). The generating parameters in panel (b) from left to right are *ψ* = 1, *ϕ* = 10^−4^, *σ*_*disp*_ = 10, *σ*_*JC*_ = 10 on the scales of 50 *×* 50, 250 *×* 250 and *ψ* = 1, *ϕ* = 10^−6^, *σ*_*disp*_ = 1, *σ*_*JC*_ = 0.1 on scales of 50 *×* 50, 250 *×* 250, respectively. The gray shading of the SAR plots represents interquantiles (minimum, 25th percentile, 75th percentile, maximum) while two quantiles (2.5th percentile, 97.5th percentiles) are added for LTT plots. In the SAD plots, the red bars represent the average number of species with abundances on log 2 scale while the blue lines represent the shape of the curve. In the spatial distribution (SPD) plots, each dot in the grid represents one individual. The color of the dot denotes the abundance-weighted mean of inverse (exponential) phylogenetic distance to the community (AWMIPD). The more red the dot is, the more phylogenetically related to the community the species of that individual is. Conversely, the blue dots represent individuals that are phylogenetically remote from the community. The yellow dots denote individuals of species with intermediate AWMIPD.

### Species abundance distribution

The SAD is affected by all four factors, i.e. *ψ, ϕ, σ*_*disp*_, *σ*_*JC*_ (Fig. 5b and Fig. S17-S25). When the individual dispersal distance is small (*σ*_*disp*_ = 0.1), there are many intermediately abundant species for the non-neutral parameter combinations (see Fig. 5b), generating a lognormal-like SAD. With such dispersal limitation, decreasing *ψ* or *ϕ* converts the species-abundance distribution from a distribution with a single interior mode (Rosindell and Cornell 2012) to a distribution with a single rare species mode of the neutral type (Rosindell et al. 2011). A large phylogenetic J-C interaction distance (*σ*_*JC*_ = 10) tends to sharpen the interior mode when dispersal is limited, indicating the species with intermediate abundance have a higher fitness. Note that under high dispersal limitation, there is always a very abundant species in the community. In contrast, a large dispersal distance (*σ*_*disp*_ = 10) produces a two-mode species-abundance distribution (Rosindell and Cornell 2012) for non-neutral parameter combinations (Fig. S17-S19). Changing the phylogenetic J-C interaction distance *σ*_*JC*_ has little influence on the shape of the species-abundance distribution. No dominant species is found in this scenario. Sampling on smaller scales does not change the abundance distribution substantially but narrows the range of abundance values. Under the protracted speciation model with large *τ* (*>* 10^5^), the number of very rare species is reduced for all scenarios. The SAD curve becomes close to the lognormal shape (Fig. S53-S61) .

### Spatial distribution

High dispersal ability of individuals tends to randomly distribute species of different phylogenetic relatedness in the community while limited dispersal ability results in patchy clustering (Fig. 5b and Fig. S26-S34). For the neutral parameter combinations, individuals with high AWMIPD dominate the community (red dots in Fig. 5b). When the dispersal distance is large, there are a few individuals with low AWMIPD distributed widely over the community (blue dots in Fig. 5b). Increasing *ψ* greatly reduces the number of phylogenetically dominant species (large AWMIPD). For the highest values of *ψ* and *ϕ*, the community is filled with individuals of distinct AWMIPD, indicating that species of different classes of phylogenetic distance to the community are distributed uniformly over the community. When the dispersal distance is small, there is a clear pattern of clustering of individuals with a large phylogenetic distance to the community for all parameter combinations. With increasing *ψ*, more low-AWMIPD individuals appear and more clusters emerge. Larger interaction distance of the phylogenetic J-C effect seems to lead to larger low-AWMIPD clusters. Decreasing the sampling scale seems to have no impact on the AWMIPD pattern.

### Lineage accumulation

The phylogenetic J-C effect tends to slow down lineage accumulation while the neutral theory yields an accelerating accumulation pattern (see the linege-through-time plots in Figs. 5b and S35-S43 and the distribution of slowdown metrics in Fig. 4 and Fig. S44-S45). The Δ*r*s for the non-neutral parameter combinations are smaller than −0.5 for all distances, suggesting a substantial slowdown in diversification. In contrast, Δ*r*s are slightly higher than 0 for the neutral model with large dispersal distance (Fig. 4). The *γ*-statistic shows a similar pattern. The Δ*r* estimates of our simulations all fall in the range of empirical estimates of the two phylogenetic data sets while the scenarios for small *ϕ* (10^−8^) and large dispersal distance (*σ*_*disp*_ = 10) and for small *ψ* (0.25) and small dispersal distance (*σ*_*disp*_ = 0.1) produce realistic *γ* estimates (Fig. S44). Further details can be found in the supplementary material (Fig. S44-S45, S48-S49). For the neutral scenarios, incorporating the protracted speciation model tends to flatten the curve, slowing down the species accumulation, which is in line with the results from the literature (Etienne and Rosindell 2012; Rosindell et al. 2015). Interestingly, protracted speciation has little influence on the curve of the scenarios with the phylogenetic J-C effect.

### Phylogenetic tree balance

Negative *β* values are generated by all parameter settings, indicating more unbalanced trees than those produced by the Yule model (Fig. S46). Phylogenetic tree balance generally increases with stronger J-C effect (larger *ψ*), and further increases with weaker phylogenetic effect (smaller *ϕ*) but the size of the effect depends on the dispersal distance and the interaction distance. A large dispersal distance tends to yield more balanced trees than a small dispersal distance. The interaction distance of the phylogenetic J-C effect seems to have little influence on the balance when the dispersal distance is large, but tends to make trees more balanced when the dispersal distance is small. The Colless index *I*_*m*_ seems to tell the same story (Fig. S47). Most trees from the two phylogenetic data sets show negative *β*-statistics and positive Colless indices, implying unbalanced trees but less imbalance than for trees generated by the neutral model (Figs. S46-S47 and Figs. S50-S51). We found no significant difference for the tree balance under the model with protracted speciation (Figs. S62-S65).

### Patterns in the BCI data

The SAR, SAD, AWMIPD and LTT plots do not differ much for the three dbh filter values (Fig. 5a). Species of three classes of AWMIPD values - low, intermediate and high values - are widely distributed over the island, which suggests a phylogenetic Janzen-Connell effect with high dispersal distance. The SAR curve shows the first two phases of the triphasic curve, being consistent with scenarios of high and intermediate dispersal distance. However, the SAD of the BCI data shows an interior mode that is a feature of the scenario with intermediate dispersal distance and low interaction. The LTT plot of the BCI data shows mild slowdowns in species accumulation at the early stage but accelerating pattern at the later stage. The Δ*r* and *γ* statistics suggest distinct species accumulation patterns: the Δ*r* statistic (-0.17) shows a slight slowdown in species accumulation, the *γ* statistic (5.36) suggests an accelerating pattern (Fig. 4), but both are higher than under any scenario except the neutral one. The Colless index and *β* statistic (*I*_*m*_ = 0.0563, *β*= −0.867) suggest a somewhat unbalanced tree shape, consistent with a strong J-C effect. Overall, the scenario of *ψ* = 1, *ϕ* = 10^−6^ with intermediate dispersal distance and low interaction distance best resembles all patterns of the BCI data (Fig. 5b).

## Discussion

We have presented a spatially explicit model based on experimental evidence that J-C effect extends to phylogenetically related species (Liu et al. 2012). In the model the J-C effect depends on the phylogenetic distance and the spatial distance between a potential colonizer and the individuals surrounding an empty site. Through simulations, we confirm the efficacy of the J-C effect to maintain high biodiversity but this is reduced by the phylogenetic relatedness effect (Fig. 3). In general, we find a palette of community patterns for different values of the strength, and the interaction distance of the phylogenetic J-C effect and the dispersal distance, ranging from low diversity in neutral scenarios to hyperdiversity of the original J-C theory (Levi et al. 2018), from a single mode (the rare species mode or the intermediately abundant species mode) to a dual mode (the rare species mode and the interior mode) species-abundance distribution (Rosindell and Cornell 2012), from pull-of-the-present lineage accumulation to slowdowns in lineage accumulation, and from unbalanced to balanced phylogenies. The SARs show a triphasic shape that is in line with previous theoretical predictions and empirical patterns (Durrett and Levin 1996; Hubbell 2001; Chave and Leigh 2002; Rosindell and Cornell 2007). However, it is attained much faster under a phylogenetic J-C effect than in the neutral scenario because the J-C effect promotes rare species and thus accumulates species faster. This supports the criticism that the dynamics of the standard neutral model are too slow to lead to empirically observed patterns within a realistic time period (Nee 2005; Ricklefs 2006; Missa et al. 2016).

The J-C effect has been shown to be pervasive and to play an important role in the maintenance of high species diversity in many communities (Petermann et al. 2008; Marhaver et al. 2013), particularly in forests (Liu et al. 2012) and grasslands (Petermann et al. 2008). Our results agree with this conclusion and demonstrate that J-C effects substantially increase species richness. We also show that extending the J-C effect to phylogenetically related species tends to reduce the diversity increase predicted by the original J-C effect alone, which only distinguished between conspecifics and heterospecifics. This implies that neglecting this phylogenetic relatedness mechanism exaggerates the power of the J-C effect to explain hyperdiversity. In practice, if closely related species share natural enemies, newly born individuals suffer negative density dependence not only from conspecifics but also from their close relatives and have a greater probability to go extinct than under a standard J-C effect with purely species-specific enemies. As a consequence, hyperdiversity cannot be maintained at the same level as by the standard J-C effect (Sedio and Ostling 2013; Stump 2017). In addition, protracted speciation reduces species richness much more strongly in the neutral scenarios than in the scenarios with the phylogenetic J-C effect. This is due to the fact that the phylogenetic J-C effect preserves rare species and hence they are on average much older than in the neutral case where they are mostly produced recently.

Furthermore, we have shown that diversity decreases with increasing dispersal, in line with the literature that argues that dispersal tends to homogenize the region and thus reduces diversity at a regional scale (Loreau 2000; Cottenie and De Meester 2004; Mouquet and Loreau 2004). This homogenization means that dominant competitors are able to distribute themselves widely and thus structure the region in a monotonous fashion (Cadotte and Fukami 2005; Cadotte et al. 2006), which directly leads to a less diverse community. In contrast, on the local community scale diversity is generally thought to be enhanced by dispersal via introducing new species (Tilman 1997; Loreau and Mouquet 1999). Our study also agrees with this conclusion: at small spatial scales larger dispersal distance leads to higher local diversity. Species are observed to distribute more uniformly as dispersal ability increases, and thus have a higher chance to explore the unfilled niches in local communities, leading to higher local diversity. In sum, the influence of dispersal on diversity differs on local and regional scales (Cadotte and Cadotte 2006), also under a phylogenetic J-C effect.

The SARs produced by the phylogenetic J-C effect show the general triphasic pattern, but the exact shape varies for different parameters of the model. This may explain the large variance in empirically observed slopes (Williamson 1988). Species with large dispersal distance tend to spread over the community, being easily sampled even when the survey area is small. This results in a fast and stable increase in the first phase, but it takes large area for the richness increase to decelerate. In contrast, a small dispersal distance reduces the sampling probability for rare species. Increasing the size of survey areas captures more rare species, which leads to a short second phase and quick arrival at the third phase. Indeed, the local, regional and continental scales of the three phases must be defined relative to the dispersal distance (Hubbell 2001; Rosindell and Cornell 2007).

Our exploration of how the phylogenetic J-C effect influences the SAD for different distances demonstrates that the J-C effect is universally disadvantageous to the most abundant species as expected but has different impact on rare or moderate species abundances depending on dispersal distance. A rare species mode arises because rare species benefit from negative density dependence. The interior mode shown in the SAD for the non-neutral parameter combinations with little dispersal limitation implies that the intermediately abundant species are not sufficiently abundant to be substantially constrained by the J-C effect but they are sufficiently abundant to be at an advantage for sampling due to high dispersal. Increasing strength of the phylogenetic J-C effect (*ψ*) decreases the number of the most abundant species and increases the number of intermediately abundant species. When the dispersal distance decreases, the intermediately abundant species benefit even more, resulting in the sharpened interior mode. This is consistent with the empirical observation of the tropical moist forest trees on Barro Colorado Island (Rosindell and Cornell 2012).

Our model predicts a broad range of species accumulation and phylogenetic balance patterns which is also observed in the empirical data (Nee et al. 1992; Mooers 1995; Mooers and Heard 1997; Blum and François 2006; Rabosky and Lovette 2008; Phillimore and Price 2008; McPeek 2008; Jønsson et al. 2012). A strong phylogenetic J-C effect tends to slow down species accumulation. Because the phylogenetic J-C effect favors rare, phylogenetically remote, species, species born in the deep past can survive, leaving a signature of an early burst of diversity. A slowdown in lineage accumulation then follows. Our model thus provides a more mechanistic explanation for the hypothesis of diversity-dependent speciation (Etienne et al. 2011). In contrast, with a weak J-C effect and/or a broad phylogenetic host range where the model tends to the neutral scenario, our model is able to account for accelerating species accumulation as follows. When reducing J-C effects, rare species are lost more quickly, leading to a high extinction rate. Consequently, rare species in the deep past went extinct but the recently born species have not had time to become extinct. This leads to the increase of the lineage accumulation rate, a phenomenon known as the pull-of-the-present (Nee et al. 1994; Phillimore and Price 2008; Rabosky and Lovette 2008). The phylogenetic J-C effect generates less imbalance than the neutral model, although abundant species are still at a sampling advantage colonizing empty sites. This is consistent with the empirical patterns. We note that the lower imbalance might result because the system may not have reached equilibrium yet. So to what extent the phylogenetic J-C effect also produces less unbalanced trees than the neutral model in equilibrium remains to be resolved. We also note that the two most extreme strengths of the phylogenetic relatedness effect that we used (*ϕ* = 0 or 1) are probably not realistic. A too high value of *ϕ* excludes the replacement of the dead individual by an individual of the same species as its neighbors. An extremely small value of *ϕ* allows natural enemies to attack any species equally. A realistic value of *ϕ* might be estimated by comparing the predicted phylogenetic tree shape and species abundance distribution with empirical data.

The comparison to the BCI data further demonstrates that our model is able to generate realistic phylogenetic and ecological patterns but a single empirical pattern can be generated by multiple scenarios with different combinations of parameters or different processes (Chave and Leigh 2002; Davies et al. 2011). Both the SAR and the AWMIPD distribution of the BCI data agree with what we observed in the scenario of high dispersal distance on the full grid scale and the scenario of intermediate dispersal distance on a local scale. The log-normal-like SAD curve is well described by the model under a strong phylogenetic Janzen-Connell effect with limited (intermediate and low) dispersal distance.This underscores a full assessment of multiple eco-evolutionary patterns to identify the underlying mechanisms (Chave et al. 2006; McGill et al. 2007). Overall, after examining all eco-evolutionary patterns that we have studied, we find that a strong phylogenetic J-C effect with intermediate dispersal distance results in patterns that are most similar to those of the BCI data.

There are some notable differences between our model predictions and the BCI patterns. The slowdown in species accumulation of the BCI data is milder than that of our simulations (but we note that our simulations are largely consistent with the two phylogenetic data sets). We see several possible reasons. First, our simulation is at a pre-equilibrium stage while the BCI data may reflect an equilibrium system. In the scenarios with high dispersal distance the original dominant species is still very abundant even under a strong phylogenetic Janzen-Connell effect. If run long enough, more rare species may appear, leading to a more realistic LTT pattern. Second, the speciation rate in our simulations is probably unrealistically high (but see Rosindell et al. 2010), which results in fast species accumulation. Third, protracted speciation (Rosindell et al. 2010; Etienne and Rosindell 2012) or random fission speciation (Etienne and Haegeman 2011) may also explain this as new species are born at higher abundance than 1. Hence, rare species are less likely to go extinct in a short time. Similarly, if tree individuals at the seedling stage are more sensitive to the phylogenetic Janzen-Connell effect (Zhu et al. 2015), one may expect that accumulating species initially takes longer time. Fourth, the phylogenetic tree for BCI may be inaccurate, particularly for the deep nodes, which may substantially affect the lineage accumulation pattern. Finally, and most importantly, the scale of the BCI 50 ha plot is much smaller than the scale at which speciation takes place and hence a local-community-level dataset such as BCI does not reflect the dynamics well (as reflected for instance by the fact that the BCI community phylogeny represents a very poorly sampled phylogeny (∼200 species) from a clade (angiosperms) with more than 300,000 species). We have attempted to take this into account by subsampling smaller areas, but this still does not represent the correct scale. Even though BCI is therefore not really appropriate to compare our model predictions, there is no other data set available yet at the regional scale for which all the patterns can be studied, and we believe that it is instructive to present the BCI analysis to illustrate our models and methods to study the interaction of ecological and evolutionary dynamics.

We emphasize that the interaction distance of the phylogenetic J-C effect and individual dispersal ability are important factors in determining species richness, spatial distribution, the SAD and phylogenetic patterns in the community. It is well-known that many species are strongly dispersal-limited (Clark and Clark 1984; Seidler and Plotkin 2006) and that dispersal limitation determines patterns of diversity and species coexistence (Macdougall and Turkington 2006; Etienne and Alonso 2007). Here we have shown that spatial phylogenetic patterns also depend crucially on dispersal. A large dispersal distance together with the phylogenetic J-C effect leads to a community where species of different phylogenetic distances distribute uniformly. In contrast, when the dispersal distance is small, a patchy community (where individuals with large phylogenetic distance to the community as a whole) is formed. This does not necessarily mean that the species within patches are phylogenetically related to each other. They can be species of low abundance and phylogenetically unrelated, which makes them equally successful in colonization. These species are less affected by the negative density effect and likely avoid natural enemies by forming such diverse subcommunities. A large interaction distance of the phylogenetic J-C effect regulates species density at a large spatial scale and consequently results in large patches, further promoting diversity. Due to computational limitations, it remains unclear whether the patchy pattern vanishes in the long run. Nevertheless, our study provides insight into patterns in a pre-equilibrium system or a transient period at the onset of community assembly. Further research analyzing the spatially explicit phylogenetic J-C effect at equilibrium and the relationship between the transient state and the equilibrium state will help in understanding the underlying mechanisms of community assembly.

Computational limitations have also barred an even more extensive exploration of parameter space, and more quantitative fitting of our model to the BCI data, using simulation-based (machine learning) techniques. We nevertheless believe that our qualitative assessment has provided a good overall match and important insights where differences between predictions and data occurred.

As we argued in the introduction, full assessment of macroecological and macroevolutionary patterns under the phylogenetic Janzen-Connell effect can complement traditional field methods to find support for the J-C effect. Here we have provided such an assessment by making predictions for a variety of patterns under a spatially explicit phylogenetic Janzen-Connell effect.

## Acknowledgments

We thank Xiaoguang Du for performing preliminary analyses with a first version of the model. We thank the Netherlands Organization (NWO) for financial support through a VICI grant awarded to RSE and the China Scholarschip Council for financial support of LX. Furthermore, we thank the Center for Information Technology of the University of Groningen for their support and for providing access to the Peregrine high performance computing cluster.

Data archival location: Once the manuscript is accepted, the data will be stored in Dryad.

